# Neutralization of SARS-CoV-2 BQ.1.1 and XBB.1.5 by Breakthrough Infection Sera from Previous and Current Waves in China

**DOI:** 10.1101/2023.02.07.527406

**Authors:** Xun Wang, Shuai Jiang, Shujun Jiang, Xiangnan Li, Jingwen Ai, Ke Lin, Shiyun Lv, Shixuan Zhang, Minghui Li, Xinyi He, Dingding Li, Chen Li, Chaoyue Zhao, Xiaoyu Zhao, Rui Qiao, Yuchen Cui, Yanjia Chen, Jiayan Li, Guonan Cai, Jixi Li, Lili Dai, Zixin Hu, Wenhong Zhang, Yanliang Zhang, Pengfei Wang

## Abstract

SARS-CoV-2 is continuing to evolve and diversify, with an array of various Omicron sub-lineages, including BA.5, BA.2.75, BN.1, BF.7, BQ.1, BQ.1.1, XBB and XBB.1.5, now circulating globally at recent time. In this study, we evaluated the neutralization sensitivity of a comprehensive panel of Omicron subvariants to sera from different clinical cohorts, including individuals who received homologous or heterologous booster vaccinations, vaccinated people who had Delta or BA.2 breakthrough infection in previous waves, and patients who had BA.5 or BF.7 breakthrough infection in the current wave in China. All the Omicron subvariants exhibited substantial neutralization evasion, with BQ.1, BQ.1.1, XBB.1, and XBB.1.5 being the strongest escaped subvariants. Sera from Omicron breakthrough infection, especially the recent BA.5 or BF.7 breakthrough infection, exhibited higher neutralizing activity against all Omicron sub-lineages, indicating the chance of BA.5 and BF.7 being entirely replaced by BQ or XBB subvariants in China in a short-term might be low. We also demonstrated that the BQ and XBB subvariants were the most resistant viruses to monoclonal antibodies. Continuing to monitor the immune escape of SARS-CoV-2 emerging variants and developing novel broad-spectrum vaccines and antibodies are still crucial.

## Introduction

It has been more than three years since the discovery of severe acute respiratory syndrome coronavirus 2 (SARS-CoV-2), the causative agent of the coronavirus disease 2019 (COVID-19) pandemic. With the global circulation and rapid evolution of the virus, five SARS-CoV-2 variants of concern (VOCs) have emerged. Among them, the Omicron variant (B.1.1.529 or BA.1) is highly divergent from the prototype virus (Wuhan-Hu-1). Moreover, since its emergence in late 2021, the Omicron variant has continued to evolve and given rise to numerous subvariants (Fig. 1a). BA.1 and its derivative BA.1.1 rapidly became dominant globally after its first detection in South Africa. After that, however, we saw a rapid surge of BA.2, which outcompeted BA.1 and became the dominant variant globally. And then, BA.2 subvariants, such as BA.2.12.1, as well as BA.4 and BA.5 became the frequently detected variants, with BA.5 gradually took the lead and turned into the most predominant variant in the world for a relatively longer period of time. Although BA.5 is still circulating with high frequency at recent time, a diverse array of Omicron sub-lineages has arisen, including BA.2.75, BN.1, BF.7, BQ.1, BQ.1.1, XBB, and more recently, XBB.1.5 (Fig. 1b). The unprecedented diversity of the so-called “variant soup” makes it harder to predict coming waves of infection^1,2^.

**Fig. 1:**
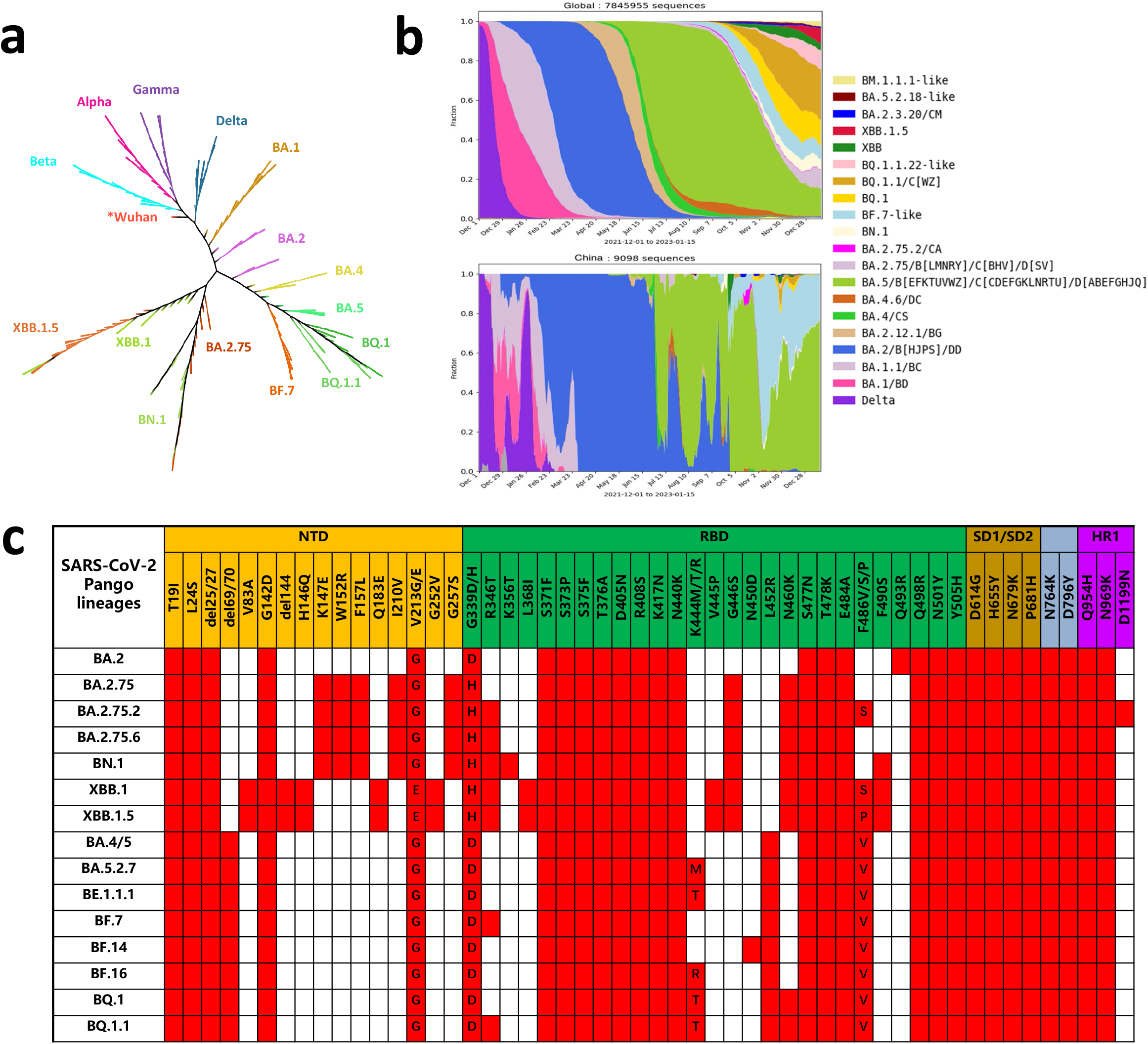
Characteristics of the Omicron subvariants. **(a)** Phylogenetic tree of the SARS-CoV-2 variants and Omicron subvariants. Twenty randomly selected sequences belonging to each of the Omicron subvariants from GISAID were used as query sequences. **(b)** Prevalence of the Omicron subvariants based on all the sequences from the globe or China available on GISAID from Dec 1^st^, 2021 to Jan 15^th^, 2023. **(c)** Spike mutations within the Omicron subvariants.

China successfully contained multiple COVID-19 outbreaks by adhering to a strict “zero-Covid” policy. However, due to the high transmissibility and immune escape properties, Omicron BA.2 caused a local outbreak in Shanghai since March 2022 and resulted in over 0.6 million laboratory-confirmed infections^3,4^. Since the “zero-Covid” policy was lifted in December 2022, China has experienced a surge in COVID-19 infections nationwide. However, the variant composition is much simpler in China, with only two main subvariants – BA.5 and BF.7 – during this current COVID-19 infection wave according to the sequences deposited in the GISAID database (Fig. 1b).

We have previously reported that SARS-CoV-2 Omicron BA.1, BA.1.1, BA.2, and BA.3 sub-lineages^5,6^, as well as BA.2 descendants (BA.2.12.1, BA.2.75, etc.), BA.4 and BA.5 (BA.4/5, share an identical spike) evaded neutralizing antibodies induced by both vaccination and infection^7^. For the recent subvariants, BF.7, BQ.1, and BQ.1.1 evolved from BA.5, with the spike protein of BF.7 having an additional R346T mutation while BQ.1 subvariant harbors K444T and N460K mutations in addition to those found in BA.5, and BQ.1.1 further obtained an R346T mutation on the top of BQ.1 (Fig. 1a and 1c). In addition, BA.2.75 carries nine additional mutations in spike compared to BA.2^7^, whereas the newly emerged BN.1 has three mutations (R346T, K356T, and F490S) in addition to those found in BA.2.75. However, the XBB and its subvariants have drawn more attention most recently. XBB resulted from a recombination between two highly diversified BA.2 lineages, with its spike containing strikingly 14 additional mutations compared to BA.2^8,9^. The XBB subvariant XBB.1 has an additional G252V mutation, while XBB.1.5 has acquired two more mutations (G252V and S486P) in the spike protein (Fig. 1c). Given the rapid growth and ever-increasing spike mutation complexity of these sub-lineages, several groups have tested the efficacy of monoclonal antibody therapeutics^8,10–14^, as well as polyclonal sera from vaccination strategies and breakthrough infections^8–10,12,14–27^ against some of these subvariant viruses. However, a comprehensive neutralization assessment of booster vaccination or breakthrough infection sera against all of these distinct emerging Omicron sub-lineages, especially an independent evaluation of the protective efficacy of sera from the previous (Delta and BA.2) and current (BA.5 and BF.7) waves in China, is still crucial for the global public health.

## Results

### Neutralization by sera from homologous or heterologous booster vaccinations

In order to evaluate the neutralization evasion properties of these distinct emerging Omicron sublineages, we constructed a panel of pseudoviruses (PsVs) representing BA.2.75, BN.1, XBB.1 XBB.1.5, BA.5, BF.7, BQ.1 and BQ.1.1, which are all circulating with high frequency at the recent time (Fig. 1b and 1c). In addition, several other Omicron subvariants with mutations of interest have also been included, such as BA.2.75.2 (BA.2.75 + R346T + F486S), BA.2.75.6 (BA.2.75 + R346T), BA.5.2.7 (BA.5 + K444M), BE.1.1.1 (BA.5 + K444T), BF.14 (BA.5 + N450D) and BF.16 (BA.5 + K444R) (Fig. 1c). And we also included wild-type (WT, D614G) and BA.2 for the sake of comparison.

We first collected serum samples from healthy adults at day 14 post homologous booster with CoronaVac, or heterologous booster with ZF2001, primed with two doses of CoronaVac (Table S1), and tested their neutralization activity on this panel of PsVs. As shown in Fig. 2a, the homologous booster group (3 × CoronaVac, n=11) had a neutralizing geometric mean titer (GMT) against WT(D614G) of 516, but the GMTs for BA.2 and BA.4/5 were 71 and 32, with 7.3- and 16-fold reductions compared to WT. However, the BA.2 and BA.5 derivative variants showed further resistance to the neutralization elicited by this booster vaccination. The BA.2 descendants, BA.2.75.2, BA.2.75.6, BN.1, XBB.1, and XBB.1.5, all had a further reduction in their neutralization titers compared to BA.2. Among them, BN.1, XBB.1 and XBB.1.5 turned out to be the most significant ones, with GMTs of 19, 18 and 13, showing substantially lower neutralization sensitivity compared with WT (27-, 29- and 40-fold reductions, respectively) and BA.2 (3.7-, 3.9- and 5.5-fold reductions, respectively). While, for the BA.5 descendants, BQ.1 and BQ.1.1 were the most resistant subvariants, with about 30-fold reduced neutralization titers compared to WT and approximately a 2-fold reduction compared to BA.5 (Fig. 2a).

**Fig. 2:**
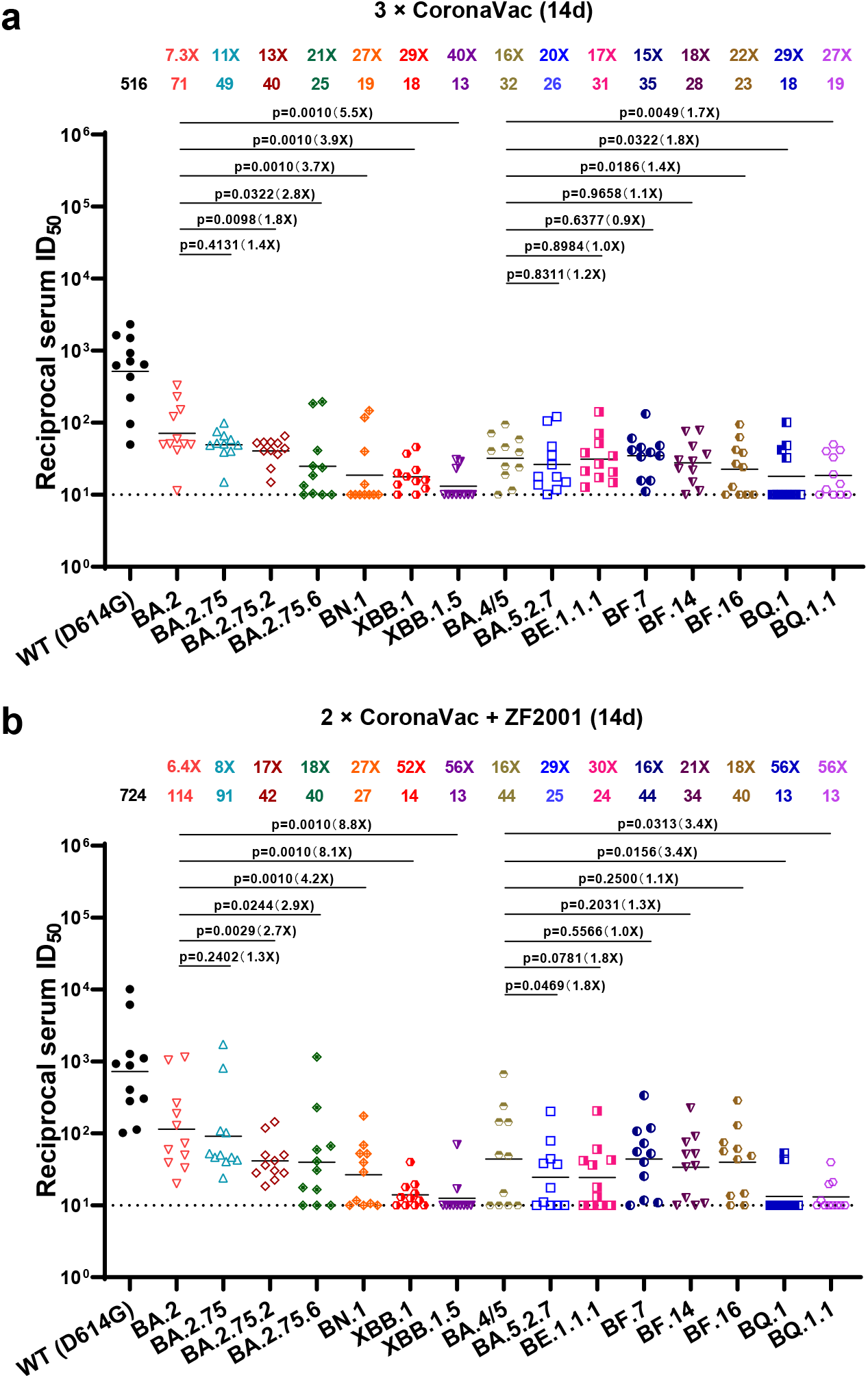
Neutralization of the Omicron subvariants by sera from homologous or heterologous booster vaccinations. Neutralization of pseudotyped WT (D614G) and Omicron sub-lineage viruses by sera collected from individuals at day 14 after vaccinated with a CoronaVac homologous booster (a) or with a ZF2001 heterologous booster dose following two doses of CoronaVac (b). For all panels, values above the symbols denote geometric mean titer and the fold-change was calculated by comparing the titer to WT. For each BA.2 or BA.5 descendant, comparison was also done with BA.2 or BA.5. *P* values were determined by using Multiple Mann-Whitney tests. The horizontal dotted line represents the limit of detection of 10. WT: wild-type.

We also observed a similar trend of neutralization resistance for this panel of Omicron sub-lineages when tested with sera collected from the heterologous booster group (2 × CoronaVac + ZF2001, n=11). Although this cohort had a higher neutralizing titer with GMTs of 724 for WT, the GMTs for BA.2 and BA.4/5 showed 6.4- and 16-fold reductions compared to WT, almost equal to the reduction levels in the homologous booster group. Similarly, the neutralizing titers against the BA.2 and BA.5 derivative variants decreased further, with again XBB.1 and XBB.1.5 to be the strongest escaped BA.2 descendants (more than 8-fold reductions compared to BA.2) and BQ.1 and BQ.1.1 being the strongest escaped BA.5 descendants (more than 3-fold reductions compared to BA.5). For these four subvariants, only a few serum samples retained marginal neutralization activity, therefore, their GMTs were almost approaching our limit of detection (LOD, 10), with reductions of strikingly >50-fold compared to that against WT (Fig. 2b). Taken together, all Omicron sub-lineages showed substantial humoral immune escape from the prototype booster vaccination, with XBB.1, XBB.1.5, BQ.1, and BQ.1.1 showing the strongest serum escape, followed by BN.1 and the other Omicron sub-lineages, in both the homologous and heterologous booster groups.

### Neutralization by breakthrough infection sera from previous waves in China

We had previously reported that BA.2 breakthrough infection significantly increased neutralizing antibodies to higher titers against BA.2, its derivative variants and BA.4/5^7^. However, those serum samples were collected and evaluated at a very early time point – day 14 post-infection, would those vaccinated individuals infected during the previous waves still maintain adequate neutralization titers against the newly emerging viruses after a longer period of time? To answer this question, we collected serum samples from individuals who had previously had Delta or BA.2 breakthrough infections at 6 months post-infection and examined the degree of neutralizing antibody escape by our panel of Omicron subvariant PsVs. For the 12 sera from Delta breakthrough infection after two inactivated vaccination doses, we noticed a trend of neutralization titer decrease for the Omicron subvariants, similar to the booster vaccination groups, and once again, BN.1, XBB.1, XBB.1.5, BQ.1 and BQ.1.1 showing the strongest serum escape, amounting to about 20-fold reductions in potency compared to WT. However, most of the samples retained detectable neutralizing activity against these distinct variants (Fig. 3a), reflecting the higher titers of the Delta breakthrough infection individuals preserved at month 6 than those of people with booster vacation only at day 14.

**Fig. 3:**
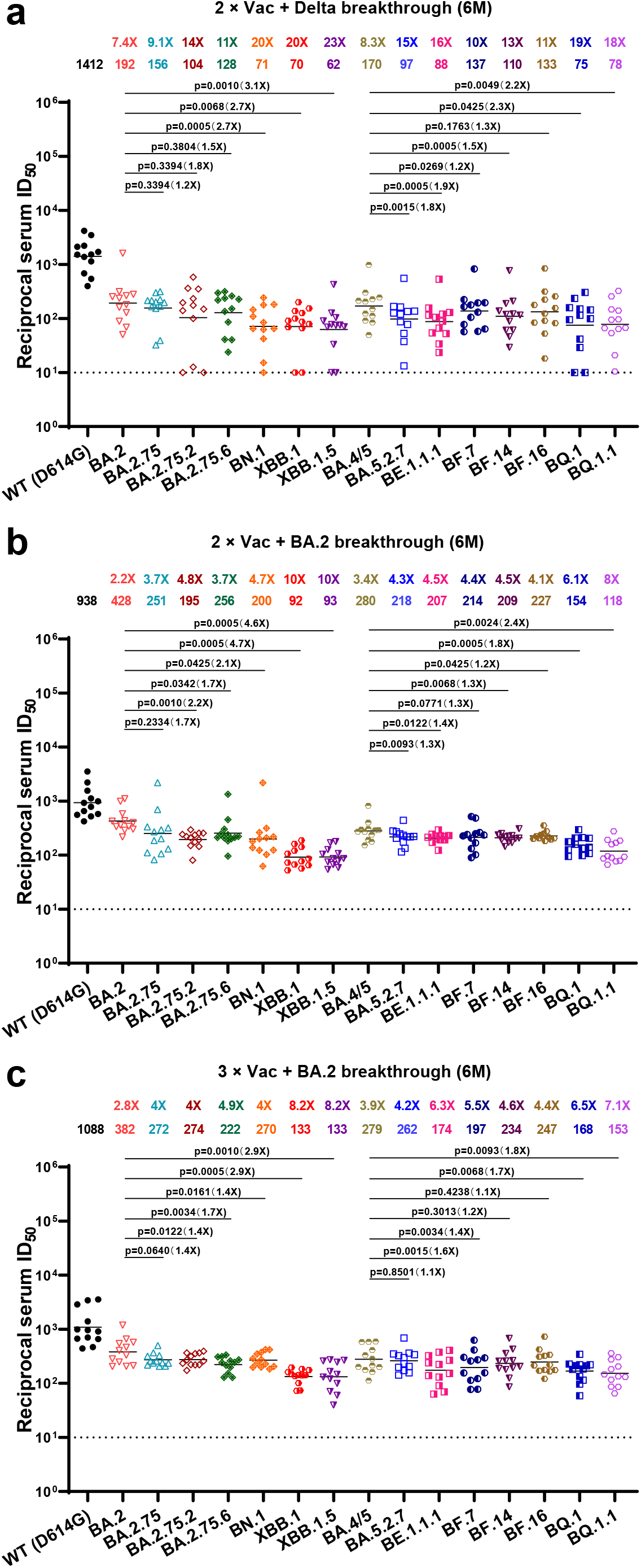
Neutralization of the Omicron subvariants by breakthrough infection sera from previous waves in China. Neutralization of pseudotyped WT (D614G) and Omicron sub-lineage viruses by sera collected from individuals at month 6 after infection with Delta virus post two doses of inactivated vaccine (a), or from those infected with BA.2 virus post two (b) or three (c) doses of inactivated vaccine in previous waves in China. For all panels, values above the symbols denote geometric mean titer and the fold-change was calculated by comparing the titer to WT. For each BA.2 or BA.5 descendant, comparisons were also done with BA.2 or BA.5. *P* values were determined by using Multiple Mann-Whitney tests. The horizontal dotted line represents the limit of detection of 10.

While for the BA.2 breakthrough infection individuals, we further divided them into two groups based on whether they had two (n = 12) or three (n = 12) inactivated vaccination doses prior to infection. In these two groups, the neutralization titers for Omicron subvariants were lower than that for WT, but the decline levels were less significant than those in the booster vaccination or Delta breakthrough infection groups. For instance, the neutralization GMTs against BA.2 dropped ~2-3-fold in BA.2 breakthrough infection groups, compared to the ~7-fold in the booster vaccination or Delta breakthrough infection groups, and similar for the other BA.2 or BA.5 descendants excluding the XBB and BQ subvariants (~4-5-fold drops in BA.2 breakthrough infection groups versus >8-fold drops in the booster vaccination or Delta breakthrough infection groups) (Fig. 2 and 3). When comparing the BA.2 breakthrough infection with the Delta breakthrough infection, with both groups had the same 2-dose vaccination history, the BA.2 breakthrough infection sera kept higher titers against BA.2 and BA.5, as well as some of their descendant viruses (Supplementary Fig. S1a), which may be associated with the antigenic difference between Omicron and Delta variants. Not surprisingly, XBB.1 and XBB.1.5 were still the strongest escaped BA.2 descendants (8-10-fold reductions compared to WT and 3-5-fold reductions compared to BA.2) and BQ.1 and BQ.1.1 were the strongest escaped BA.5 descendants (6-8-fold reductions compared to WT and ~2-fold reductions compared to BA.5) (Fig. 3b and 3c). However, it is noteworthy that even for these strongest escaped viruses, all of the samples from BA.2 breakthrough infection post two or three vaccination doses retained detectable neutralizing activity, in contrast to what we observed for booster vaccination-only sera.

### Neutralization by breakthrough infection sera from the current wave in China

We then examined the resistance of these Omicron subvariants to serum samples from individuals who had more recently had BA.5 or BF.7 breakthrough infections. For the 20 BA.5 breakthrough infection individuals, 9 participants had received two-dose inactivated vaccines (2 × Vac + BA.5 breakthrough) and 11 had homologous booster (3 × Vac + BA.5 breakthrough), and serum samples were collected at day 14 post breakthrough infections. Overall, BA.5 breakthrough infection induced pretty high neutralizing titers for WT and Omicron subvariants. The GMTs for BA.5 and some of its derivative variants like BF.7 were all above 1000, and even for the BQ.1 and BQ.1.1 subvariants, the GMTs were sustained above 300. Certain BA.2 descendant viruses, such as BA.2.75.2, BN.1, XBB.1 and XBB.1.5 exhibited stronger evasion from the BA.5 breakthrough infection sera. But even the neutralization titers for XBB.1 and XBB.1.5 decreased 9-14-fold compared to WT and 5-7-fold compared to BA.2, they still retained GMTs ~200 (Fig. 4a and 4b). Moreover, similar to the BA.2 breakthrough infection, the BA.5 breakthrough infection elicited similar levels of neutralizing antibodies in vaccinated people regardless of whether they had booster (Supplementary Fig. S1b and S1c).

**Fig. 4:**
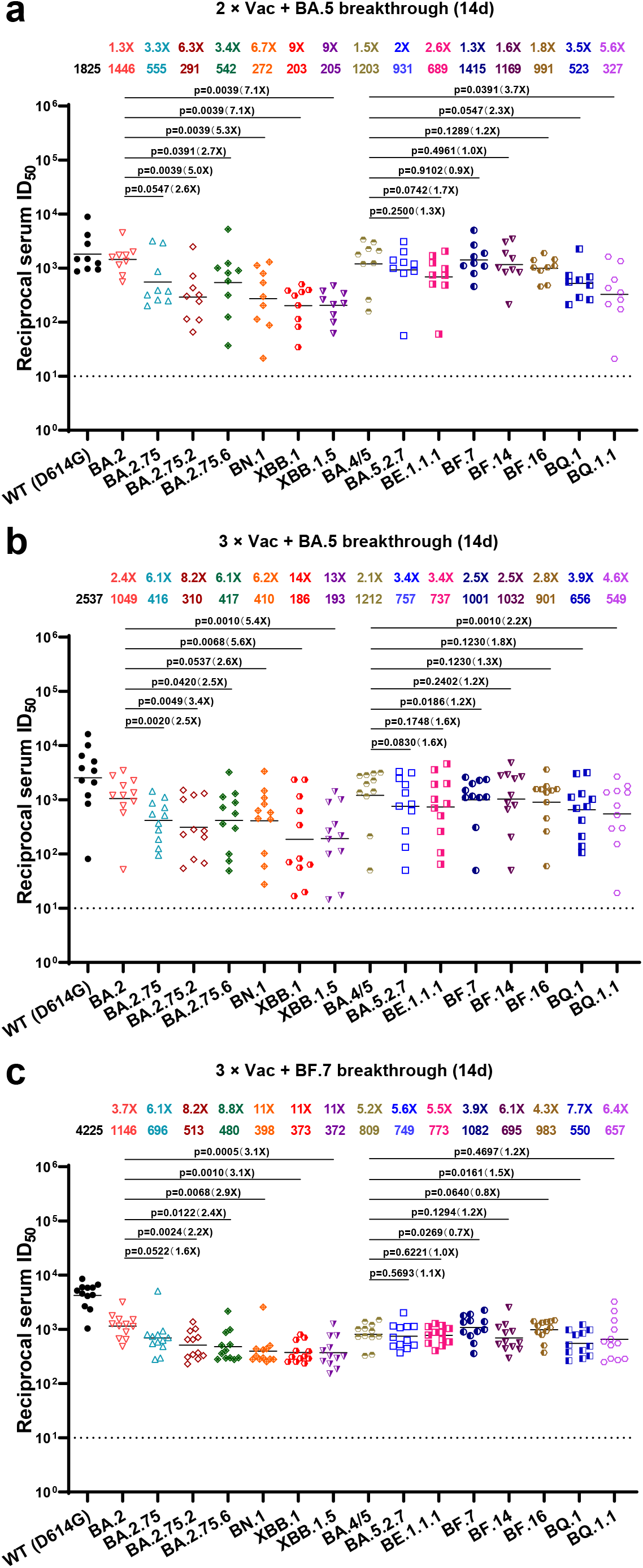
Neutralization of the Omicron subvariants by breakthrough infection sera from current wave in China. Neutralization of pseudotyped WT (D614G) and Omicron sub-lineage viruses by sera collected from individuals at day 14 after infection with BA.5 virus post two (a) or three (b) doses of inactivated vaccine, or from those infected with BF.7 virus post three doses of inactivated vaccine (c) in the current wave in China. For all panels, values above the symbols denote geometric mean titer and the fold-change was calculated by comparing the titer to WT. For each BA.2 or BA.5 descendant, comparisons were also done with BA.2 or BA.5. *P* values were determined by using Multiple Mann-Whitney tests. The horizontal dotted line represents the limit of detection of 10.

For the BF.7 breakthrough infection group, we have a cohort of 12 people, all had received three-dose inactivated vaccines prior to infection. All participants had relatively high neutralization activities against BA.5 and its derivative variants, including BQ.1 and BQ.1.1 (GMTs >500), with the highest titer against BF.7 virus as expected. Although some BA.2 descendants, BN.1, XBB.1 and XBB.1.5 exhibited the lowest neutralization sensitivity, which might be explained by their antigenic distance from BF.7, their neutralization GMTs were all above 300 (Fig. 4c). When compared with the above 3 × Vac + BA.5 breakthrough infection data, the BF.7 breakthrough infection mounted generally comparable neutralizing responses (Supplementary Fig. S1d). Taken together, these results indicated that people vaccinated and then infected in the current wave in China should have a good chance of being protected from the currently circulating SARS-CoV-2 viruses, at least for a short period of time.

### Summary of neutralization by polyclonal sera

To have a more direct parallel comparison of serum-neutralizing activity from individuals with various vaccination/infection statuses, we integrated data from Fig. 2–4 and summarized in Fig. 5. Since the BA.2 or BA.5 breakthrough infection showed no apparent difference regarding whether they had received two or three prior vaccination doses (Supplementary Fig. S1b and S1c), we combined the “2 × Vac + BA.2 breakthrough infection” and “3 × Vac + BA.2 breakthrough infection” groups into a “BA.2 breakthrough infection” group, and did the same thing for the “BA.5 breakthrough infection” group. As shown in Fig. 5a, people with booster vaccination only had the lowest serum neutralization activity, with almost undetectable titers against BQ.1, BQ.1.1, XBB.1, and XBB.1.5. Those vaccinated and infected by Delta or BA.2 variants in previous waves still retained appreciable activity against the distinct Omicron subvariants 6 months post-infection. The highest neutralization activity came from those vaccinated and infected with BA.5 or BF.7 viruses in the current wave in China, which could be attributed to the higher antigenic similarity of BA.5 or BF.7 with the newly emerged viruses as well as the shorter sample collection time post-infection. Large proportion of the sera from the booster vaccination-only groups had neutralizing antibody titers under our LOD, especially for BQ.1, BQ.1.1, XBB.1, and XBB.1.5 subvariants. However, all sera from the Omicron breakthrough infection groups had neutralizing antibody titers above LOD, and the overall trend of curves shifting to the right further indicated the enhanced neutralization potency (Fig. 5b).

**Fig. 5:**
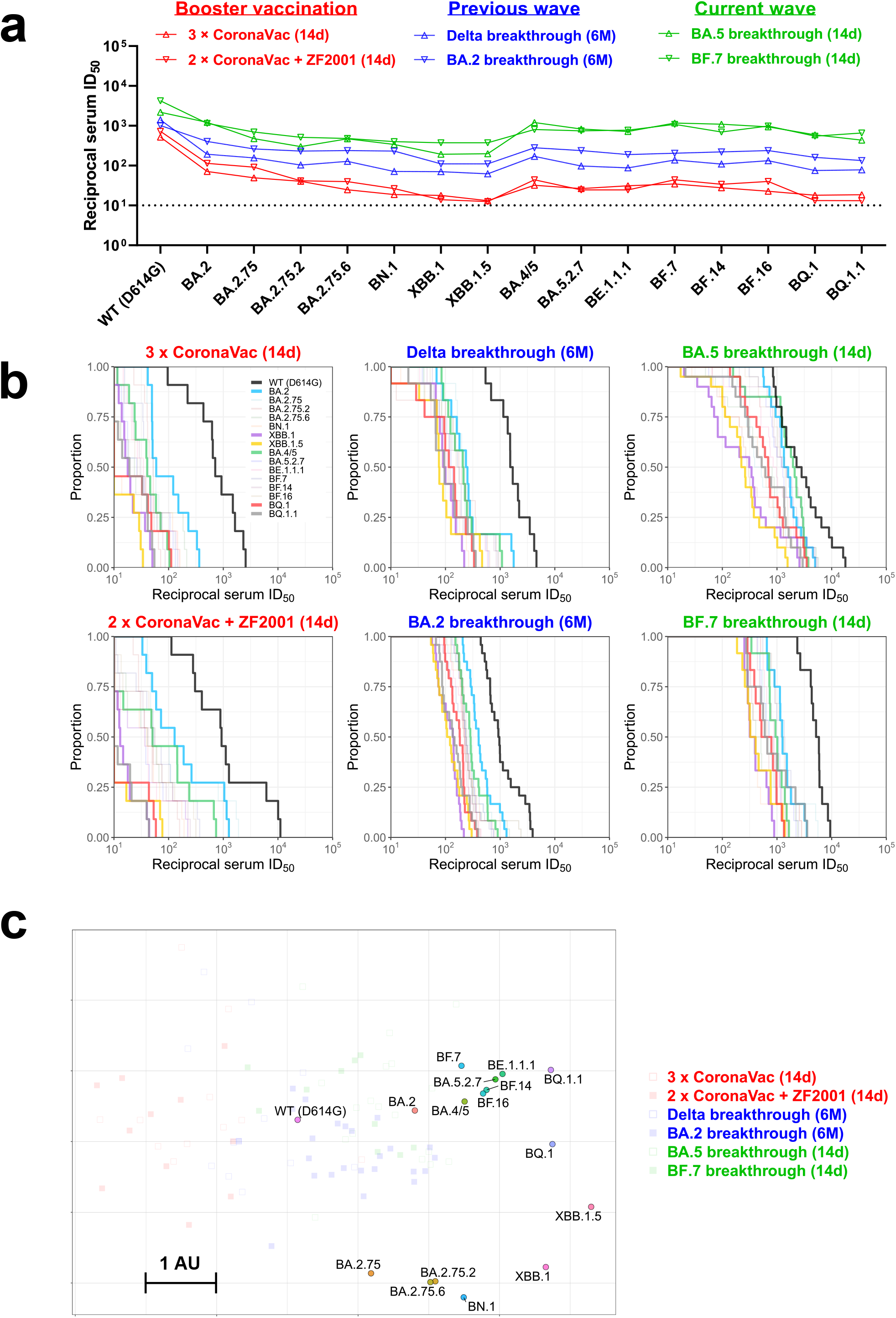
Summary of neutralization by polyclonal sera. **(a)** In parallel comparison of neutralization GMTs against distinct Omicron subvariants by sera collected from individuals with homologous or heterologous booster vaccinations, or with breakthrough infections in the previous or current waves in China. **(b)** Cumulative distribution function plots of titers against WT (D614G) and Omicron subvariants (with BA.2, BA.4/5, BQ.1, BQ.1.1, XBB.1, and XBB.1.5 highlighted), showing the proportion of samples at or above a given titer. **(c)** Antigenic map based on the serum neutralization data from Fig. 2–4. Virus positions are represented by closed circles whereas serum positions are shown as open or closed squares. Both axes represent antigenic distance with one antigenic distance unit (AU) in any direction corresponding to a 2-fold change in neutralization ID_50_ titer.

To visualize and quantify the antigenic distances among WT and the Omicron subvariants, we utilized all the serum neutralization results to construct an antigenic map (Fig. 5c). The map reveals features of the antigenic evolution of the SARS-CoV-2 virus. While some Omicron subvariants like BA4/5, BA.5.2.7, BE.1.1.1, BF.14, and BF.16 tend to group in a cluster, the BQ and XBB subvariants have drifted away from BA.2 and BA.4/5 antigenically, with XBB.1.5 to be the most antigenically distinct subvariants. Based on each antigenic unit equaling a 2-fold difference in virus neutralization, BQ.1 and BQ.1.1 are approximately 12-fold more resistant to serum neutralization than the ancestral D614G, but only about 2.5-fold more resistant than its predecessor BA.5. While XBB.1.5 is over 20-fold more resistant to serum neutralization than D614G, 7-fold more resistant than its predecessor BA.2, and 5-fold more resistant than BA.5, raising concerns about how the antigenic drift will affect vaccine efficacy in the real world.

### Neutralization by monoclonal antibodies

For individuals at high risk, for example, those who are immunocompromised, monoclonal antibodies (mAbs) or cocktails of mAbs are administered as prophylaxis or therapy. Therefore, it’s crucial to evaluate the neutralizing activity of mAbs against the emerging Omicron subvariants. Susceptibility of our panel of viruses to neutralization by 12 mAbs targeting the spike and 2 mAbs targeting the human angiotensin converting enzyme-2 (ACE-2) receptor was tested. The neutralization IC_50_ values are displayed in Fig. 6a and the mAbs’ fold changes in IC_50_ compared to WT are shown in Fig. 6b. LY-CoV1404 (bebtelovimab)^28^, the only clinically authorized/approved mAb kept its potent neutralization activity against BA.1, BA.2 and BA.3 in our previous study^5^, was totally inactive against BE.1.1.1, XBB.1, XBB.1.5, BQ.1 and BQ.1.1, probably due to the K444T and V445P mutations^8^. Interestingly, the different K444 mutations exhibited different sensitivity to LY-CoV1404, which retained neutralizing activity against BA.5 (no K444 mutation) and BF.16 (BA.5 + K444R), lost activity partially against BA.5.2.7 (BA.5 + K444M), but completely lost activity against BE.1.1.1 (BA.5 + K444T). Structural analysis indicated that the K444R mutation maintained the hydrogen bonds linking spike K444 to D57 and D58 of the antibody heavy chain. While both the K444M and K444T mutations influenced the distance and number of the hydrogen bonds, K444M could form another hydrogen bond with Y54 of the antibody. The docking energy of LY-CoV1404 heavy chain to K444T was predicted to be higher than that to K444M as well (Fig. 6c), consistent with the neutralization results. S309 (sotrovimab)^29^, with emergency use authorization (EUA) granted but withdrawn later, lost its neutralization activity partially but retained detectable activity against all these Omicron subvariants. Brii-196 (amubarvimab)^30^, which has been approved in China in combination with romlusevimab, although kept low activity against some subvariants like BA.5 and BF.7, were rendered inactive against BA.2.75.2, XBB.1, XBB.1.5, BQ.1 and BQ.1.1. We also tested COV2-2130 (cilgavimab) and COV2-2196 (tixagevimab)^31^, as well as their marketed combination (Evusheld). Despite BA.2 and BA.5 showing reduced but retained sensitivity to COV2-2130 and Evusheld, most of the other subvariants were entirely resistant to COV2-2130, COV2-2196, and their combination.

**Fig. 6:**
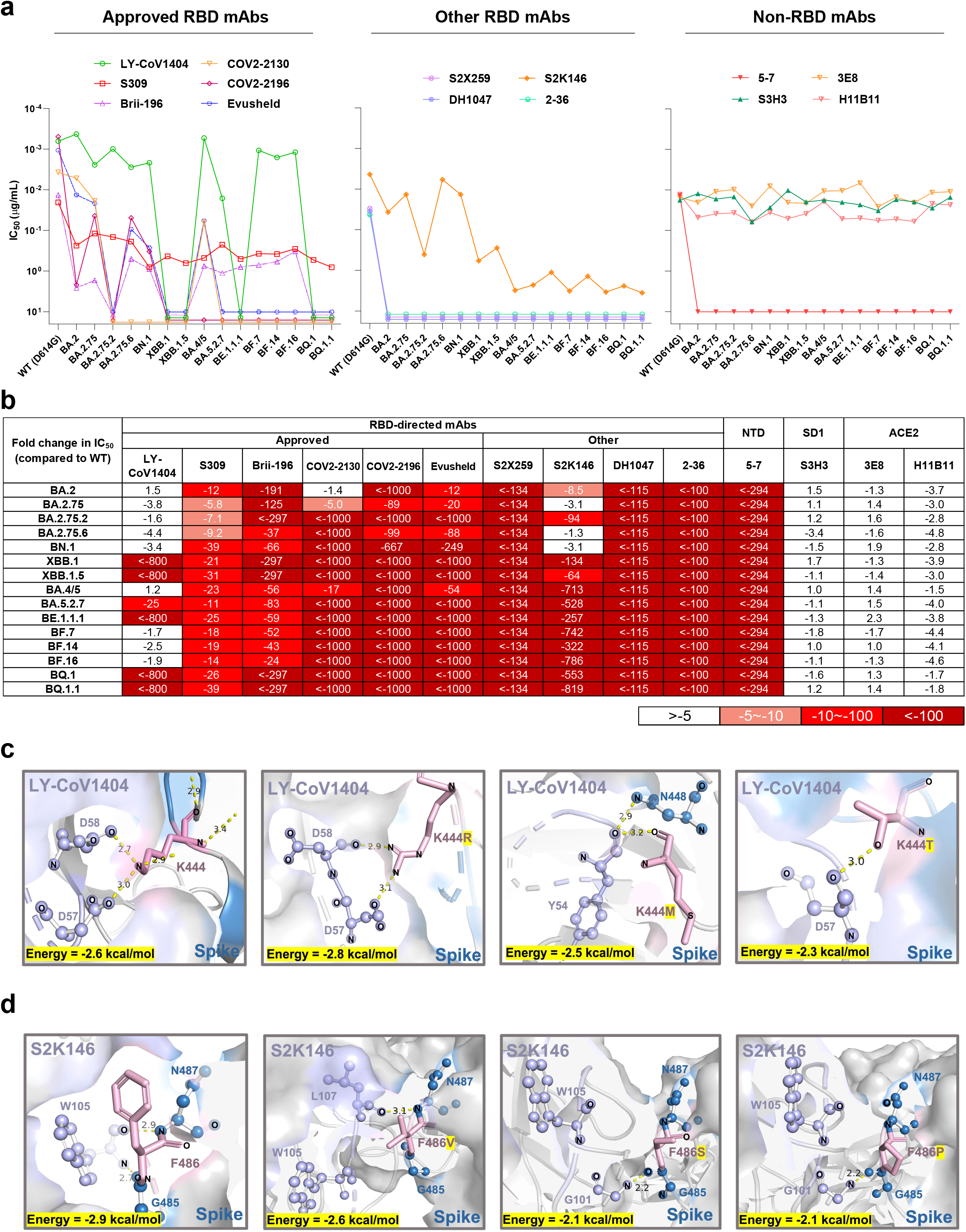
Neutralization of the Omicron subvariants by mAbs. **(a)** Neutralization of pseudotyped WT (D614G) and Omicron sub-lineage viruses by mAbs targeting different epitopes. Changes in neutralization IC_50_ are shown. **(b)** Fold increase or decrease in neutralization IC_50_ of mAbs against Omicron sub-lineage pseudoviruses relative to WT, presented as a heat map with darker colors implying greater change. **(c)** Structural modeling analysis of K444R/M/T effects on binding of LY-CoV1404. **(d)** Structural modeling analysis of F486V/S/P effects on binding of S2K146.

We next evaluated some other RBD-directed mAbs of interest, including S2X259^32^, DH1047^33^, S2K146^34^, and 2-36^35,36^, all have been reported with some extent of broadly neutralizing activity. Unfortunately, S2X259, DH1047, and 2-36 lost their activity against all the Omicron subvariants. S2K146 retained activity against BA.2, BA.2.75, BA.2.75.6, and BN.1, but lost activity significantly against all the other Omicron subvariants, well correlated with whether these viruses contain F486 mutation or not (Fig. 1c and 6a). Structural modeling revealed that the F486V/S/P mutations could destroy the original hydrogen bonds linking spike G485 and N487 to W105 on S2K146 heavy chain. Also, the predicted docking energy of the F486 mutations increased, making S2K146 more likely to dissociate from the viral spike (Fig. 6d). We also tested four more non-RBD mAbs, including 5-7 targeting the NTD^35,37^, and S3H3 targeting SD1^38^, as well as two anti-ACE2 antibodies, 3E8^39^, and H11B11^40^. Although 5-7 was reported previously^5^ to be active against BA.1 and BA.3, it lost activity completely against the BA.2 and BA.5 descendants. However, the silver lining is that S3H3, 3E8 and H11B11 kept their neutralization potency against all the current Omicron subvariants with little change (Fig. 6a and 6b), indicating that outside of the mutation hotspots, there are still regions on the viral spike and receptor that could be targeted to race against viral evolution.

## Discussion

Although the worst days of the COVID-19 pandemic, which has claimed the lives of at least 6.5 million individuals worldwide, may be behind us, novel SARS-CoV-2 variants will likely continue to present considerable public health challenges around the globe for years^41^. The continued evolution of the virus has led to the emergence of the Omicron variant and numerous sub-lineages, requiring continued warnings and evaluations. Currently, the global circulation of the virus is made up of a diverse array of Omicron sub-lineages, including BA.5, BA.2.75, BN.1, BF.7, BQ.1, BQ.1.1, XBB, and XBB.1.5, etc. (Fig. 1b). With the rapid rise and an increasing number of mutations, the emerging Omicron subvariants are predicted to obtain enhanced immune evasion to current vaccines and mAb therapeutics.

In this study, we first created a comprehensive panel of Omicron subvariant pseudoviruses and evaluated their neutralization sensitivity to sera from eight different clinical cohorts, including individuals who received homologous or heterologous booster vaccinations, patients who had Delta or BA.2 breakthrough infection after vaccination in previous waves, and patients who had BA.5 or BF.7 breakthrough infection in the current wave in China (Fig. 2–4). Overall, all the Omicron subvariants exhibited substantial neutralization evasion compared to WT, with XBB subvariants to be the strongest escaped BA.2 descendants and BQ subvariants to be the strongest escaped BA.5 descendants, which is consistent with some recent studies^10,42^. However, we included more viruses here, including the most recently discovered XBB.1.5, which is rapidly spreading in the USA. We found that XBB.1.5 did not exhibit enhanced neutralization resistance over its parental XBB.1; therefore, its enhanced transmissibility might be due to other reasons like the greater ACE2 binding^17,19^. Sera from individuals who received prototype vaccine boosters and those with Delta breakthrough infections, lost greater neutralizing sensitivity to Omicron sub-lineages than those with Omicron breakthrough infections, which could be attributed to the big antigenic difference between Omicron and pre-Omicron viruses. In addition, for people with Omicron breakthrough infections, a previous booster vaccination did not elicit higher neutralizing activity. Gao et al. also reported that in patients with BA.2 breakthrough infection, additional doses of prototype vaccine might recall a robust immune response to target the original strain, but dampen immune responses to newly infected strains^43^. When compared in parallel (Fig. 5), sera from the recent BA.5 or BF.7 breakthrough infections exhibited the highest neutralizing activity against all Omicron sub-lineages, indicating that people vaccinated previously and infected in the current wave in China should have a certain degree of protection against re-infection by the emerging Omicron subvariants including BQ and XBB. Although further studies are needed to see how long this hybrid immunity could maintain, given the high infection rate in the current wave, the chance of BA.5 and BF.7 being completely replaced by other viruses in China in a short-term might be low.

We also showed that the panel of Omicron subvariants were entirely or partially resistant to neutralization by most mAbs tested, including the currently approved bebtelovimab, amubarvimab and Evusheld. Our data were in line with observations by others^11,14,42^, but with extra detailed impacts of the K444R/M/T and F486V/S/P mutations on antibody binding. Consistent with the polyclonal sera results, BQ and XBB subvariants also became the most resistant viruses to mAbs. S3H3, targeting SD1, as well as 3E8 and H11B11, targeting ACE2, retained neutralizing activity against all the Omicron subvariants. It would be interesting to investigate further to determine whether these mAbs could provide benefits in clinical use.

Collectively, the SARS-CoV-2 virus continues to evolve and increase immune evasion to vaccination, breakthrough infection, and mAb treatment. The significantly higher neutralizing titers elicited by Omicron breakthrough infections than prototype vaccination boosters suggested developing novel vaccines based on emerging variants to better control the forthcoming viruses. Universal strategies applicable to all SARS-CoV-2 variants and other emerging zoonotic coronaviruses, such as those based on the conserved region of spike or the ACE2-based therapeutics^44^, should be explored. Continued vigilance and sustained support for development of updated vaccine and mAb therapies that protect broadly are also of great importance.

## DATA AVAILABILITY

All the data are provided in the main or the supplementary figures.

## ACKNOWLEDGMENTS

This study was supported by funding from the National Natural Science Foundation of China (32270142 to P.W., 82000070 to Shuai Jiang, 82272320 to L.D.), Shanghai Rising-Star Program (22QA1408800 to P.W.), Nanjing Research Center for Infectious Diseases of Integrated Traditional Chinese and Western Medicine (YBZX2022 to Y.Z.), Shanghai Municipal Science and Technology Major Project (HS2021SHZX001 to W.Z.), Shanghai Science and Technology Committee (21NL2600100 and 20dz2260100 to W.Z.), Beijing Natural Science Foundation (7222092 to L.D.), National Key Research and Development Project of China (2021YFC2301500 to J.L.), and was also partly supported by a grant from the major project of Study on Pathogenesis and Epidemic Prevention Technology System (2021YFC2302500) by the Ministry of Science and Technology of China.

## AUTHOR CONTRIBUTIONS

P.W., Y.Z., W.Z., Z.H., L.D., and J.L. designed and supervised the study; X.W., Shuai J., Shujun J., X.L., J.A., K.L., S.L., S.Z., and M.L. performed the experiments with help from X.H., C.L., C.Z., X.Z., R.Q., Y. Cui, Y. Chen., J.L., and G.C.; Shujun J., J.A., K.L., S.L., and D.L. provided critical materials; P.W., Y.Z., W.Z., Z.H., L.D., J.L., X.W., Shuai J., Shujun J., X.L., J.A., K.L., S.L., S.Z., and M.L. analyzed the data and wrote the manuscript. All authors reviewed, commented, and approved the manuscript.

## Materials and methods

### Serum samples

Sera from individuals who received two or three doses of inactivated vaccine (CoronaVac) or recombinant protein subunit (ZF2001) vaccine were collected at Nanjing Hospital of Chinese Medicine 14 days after the final dose. 12 individuals who were breakthrough infected with SARS-CoV-2 Delta (B.1.617.2) variant after receiving two doses of inactivated vaccine were recruited at the Nanjing Hospital of Chinese Medicine. 24 individuals who were infected with BA.2 variant after receiving two or three doses of inactivated vaccine were recruited at Huashan Hospital, Fudan University. 20 individuals who were infected with BA.5 variant after receiving two or three doses of inactivated vaccine were recruited at the Nanjing Hospital of Chinese Medicine. 12 individuals who were infected with SARS-CoV-2 BF.7 variant after receiving three doses of inactivated vaccine were recruited at the Youan Hospital, Capital Medical University. The SARS-CoV-2 infection of all the subject was confirmed by sequencing. Their baseline characteristics are summarized in Table S1. All the participants provided written informed consents. All collections were conducted according to the guidelines of the Declaration of Helsinki and approved by the ethical committee of Huashan Hospital Affiliated to Fudan University (number KY2022-596 & KY2021-749) and Nanjing Hospital of Chinese Medicine Affiliated to Nanjing University of Chinese Medicine (number KY2021162).

### Cell lines

Expi293F cells (Thermo Fisher Cat# A14527) were cultured in the serum free SMM 293-TI medium (Sino Biological Inc.) at 37 °C with 8% CO_2_ on an orbital shaker platform. HEK293T cells (Cat# CRL-3216), Vero E6 cells (cat# CRL-1586) were from ATCC and cultured in 10% Fetal Bovine Serum (FBS, GIBCO cat# 16140071) supplemented Dulbecco’s Modified Eagle Medium (DMEM, ATCC cat# 30-2002) at 37°C, 5% CO_2_. I1 mouse hybridoma cells (ATCC, cat# CRL-2700) were cultured in Eagle’s Minimum Essential Medium (EMEM, ATCC cat# 30-2003)) with 20% FBS.

### Monoclonal antibodies

Monoclonal antibodies tested in this study were constructed and produced at Fudan University. For each antibody, variable genes were optimized for human cell expression and synthesized by HuaGene™ (China). VH and VL were inserted separately into plasmids (gWiz or pcDNA3.4) that encode the constant region for H chain and L chain. Monoclonal antibodies were expressed in Expi293 (ThermoFisher, A14527) by co-transfection of H chain and L chain expressing plasmids using Polyethylenimine and culture at 37 °C with shaking at 125 rpm and 8% CO_2_. On day 5, antibodies were purified using MabSelect™ PrismA (Cytiva, 17549801) affinity chromatography.

### Construction and production of variant pseudoviruses

Plasmids encoding the WT (D614G) SARS-CoV-2 spike and Omicron sub-lineage spikes were constructed. HEK293T cells were transfection with the indicated spike gene using Polyethylenimine (Polyscience). Cells were cultured overnight at 37°C with 5% CO_2_ and VSV-G pseudo-typed ΔG-luciferase (G*ΔG-luciferase, Kerafast) was used to infect the cells in DMEM at a multiplicity of infection of 5 for 4 h before washing the cells with 1×DPBS three times. The next day, the transfection supernatant was collected and clarified by centrifugation at 3000g for 10 min. Each viral stock was then incubated with 20% I1 hybridoma (anti-VSV-G; ATCC, CRL-2700) supernatant for 1 h at 37 °C to neutralize the contaminating VSV-G pseudotyped ΔG-luciferase virus before measuring titers and making aliquots to be stored at −80 °C.

### Pseudovirus neutralization assays

Neutralization assays were performed by incubating pseudoviruses with serial dilutions of monoclonal antibodies or sera, and scored by the reduction in luciferase gene expression. In brief, Vero E6 cells were seeded in a 96-well plate at a concentration of 2×10^4^ cells per well. Pseudoviruses were incubated the next day with serial dilutions of the test samples in triplicate for 30 min at 37 °C. The mixture was added to cultured cells and incubated for an additional 24 h. The luminescence was measured by Luciferase Assay System (Beyotime). IC_50_ was defined as the dilution at which the relative light units were reduced by 50% compared with the virus control wells (virus + cells) after subtraction of the background in the control groups with cells only. The IC_50_ values were calculated using nonlinear regression in GraphPad Prism.

### Antigenic Cartography

The constructed antigenic map was based on serum neutralization data utilizing the antigenic cartography methods^45,46^, which are implemented in the Racmacs package (https://acorg.github.io/Racmacs/). The antigenic map was generated in R with 10000 optimization steps and other default parameters in a 2-dimesional space. The distances between positions of sublineages and serum on the antigenic map were optimized so that distances approach the fold decreases in neutralizing ID_50_ titer, relative to the maximum titer for each serum. Each unit of distance in arbitrary directions in the antigenic map represents a 2-fold change in the ID_50_ titer.

### ID_50_ Cumulative Distribution Analysis

The ID_50_ cumulative distributions of different sub-lineages were determined by the proportion of samples at or above a given titer at different vaccination/infection statuses. The max ID_50_ on the Cumulative Distribution figure were assumed as the 1.1-fold of the maximum titer for each sublineage.

### Prevalence of Different Sub-lineages

The prevalences of different Omicron sub-lineages were summarized using Embers in COVID-19 Viral Genomic Analysis pipeline^47^ (https://cov.lanl.gov/content/index), whose data were from GISAID. The prevalence data for Global and China were collected from Dec 1^st^, 2021 to Jan 15^th^, 2023 and summarized by “Grouped Pango lineages”.

### Sequence alignment and phylogenetic tree construction

This analysis involved 320 nucleotide sequences, including 20 samples for each lineage (Wuhan, Alpha, Beta, Gamma, Delta, BA.1, BA.2, BA.4, BA.5, BA.2.75, BN.1, BF.7, BQ.1, BQ.1.1, XBB.1, XBB.1.5) randomly selected from GISAID database. Sample set can be found at https://github.com/wenrurumon/GISAID/blob/main/2022.04.07.487489/map2.csv. Sequence alignment was carried out using MAFFT progress ^48^ and corrected manually. The evolutionary history was inferred using the Neighbor-Joining method. In order to present the evolutionary relationship more accurately and finely, we adjusted the samples. There was a total of 155 sequences in the final dataset. The optimal tree was shown. Evolutionary analysis was conducted in FastTree ^49^ and visualized in iTOL v6.

### Antibody binding and mutagenesis analysis

All the structures were downloaded from the PDB (7MMO for LY-CoV1404 and 7TAS for S2K146) for analysis. The interface residues were obtained by running the Interface Residues script from PyMOLWiki in PyMOL. Site-directed mutagenesis was also conducted in PyMOL. The molecular docking section uses Autodock Vina, and Autodock Grid Box to define the active centers (K444 and F486) and add polar hydrogen atoms and Kollman charges. The results were used to evaluate the effect of mutation of the spike on antibody binding ability using binding energy (affinity energy/(kcal/mol)) and hydrogen-kin distance (nm) and its quantity. Active regions include F486 of the spike (sequence: ‘EGFNCYF’, 7TAS chain E), K444 of the spike (sequence: ‘EGFNCYF’, 7MMO chain F), LY-CoV1404 (sequence: ‘YWDDDKR’, 7MMO chain D), S2K146 (sequence: ‘DLGRGGWYL’, 7TAS chain H). All structural analysis maps were generated in PyMOL v.2.5.4 (Schrödinger, LLC).

### Quantitative and statistical analysis

The statistical analyses for the pseudovirus virus neutralization assessments were performed using GraphPad Prism for calculation of mean value for each data point. Each specimen was tested in triplicate. Antibody neutralization IC_50_ values were calculated using a five-parameter dose-response curve in GraphPad Prism. For comparing the serum neutralization titers, statistical analysis was performed using Multiple Mann-Whitney tests. Two-tailed p values are reported. No statistical methods were used to determine whether the data met assumptions of the statistical approach.

