## Supplementary information for "Neutralization of SARS-CoV-2 BQ.1.1 and XBB.1.5 by Breakthrough Infection Sera from Previous and Current Waves in China"

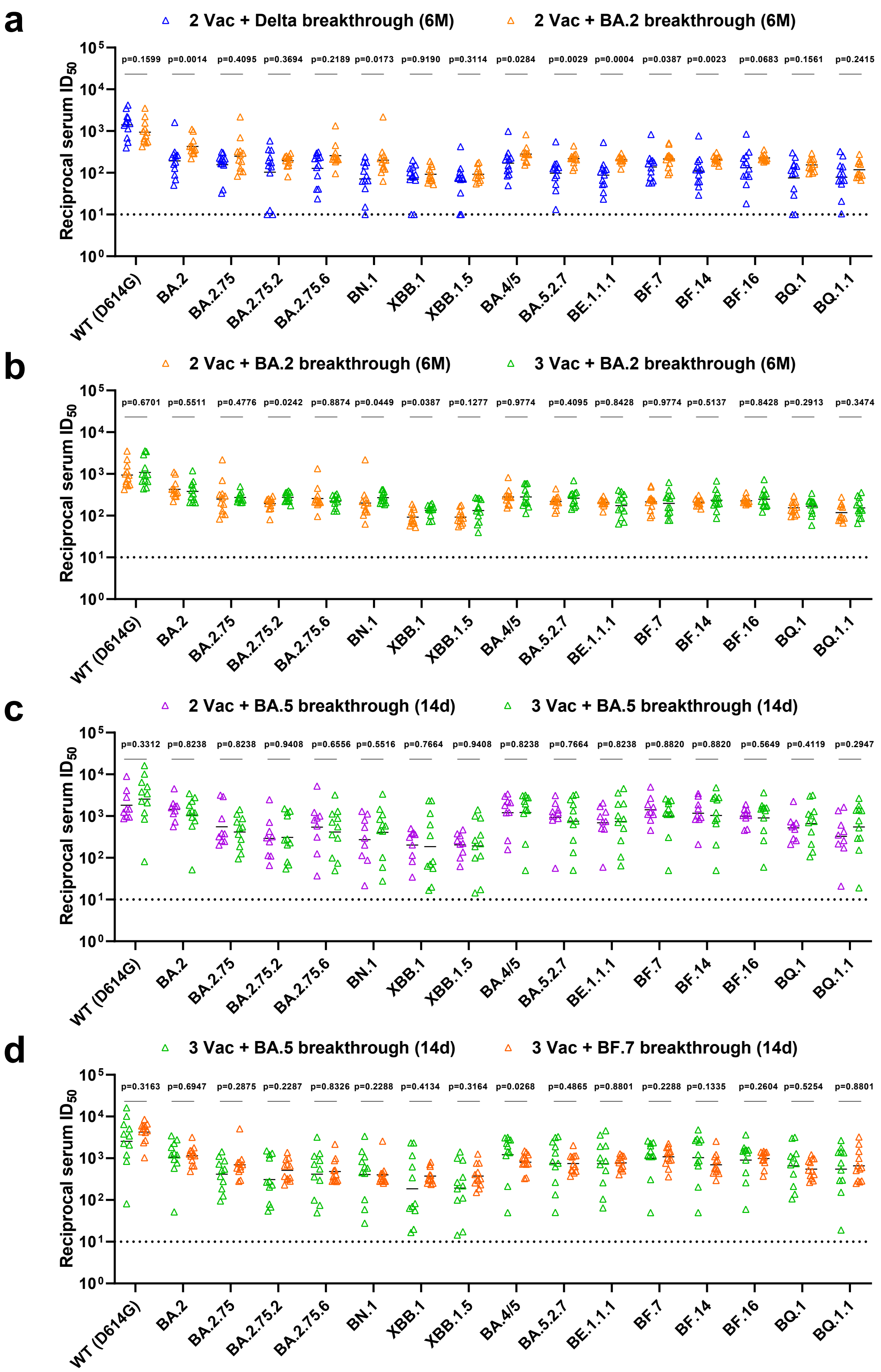


**Supplementary Fig. S1. In parallel comparison of serum neutralization titers against distinct SARS-CoV-2 variants.** (a) Neutralization titers of sera collected at month 6 from individuals with

Delta breakthrough infection versus those with BA.2 breakthrough infection after 2 doses of inactivated vaccinations. (b) Neutralization titers of sera collected at month 6 from individuals with BA.2 breakthrough infection after 2 doses versus those after 3 doses of inactivated vaccinations. (c) Neutralization titers of sera collected at day 14 from individuals with BA.5 breakthrough infection after 2 doses versus those after 3 doses of inactivated vaccinations. (d) Neutralization titers of sera collected at day 14 from individuals with BA.5 breakthrough infection versus those with BF.7 breakthrough infection after 3 doses of inactivated vaccinations. *P* values were determined by multiple Mann-Whitney tests.

### Supplementary Table S1. Baseline characteristics of enrolled participants

| | 3 × CoronaVac (n=11) | 2 × CoronaVac + ZF2001 (n=11) | 2 × Vac + Delta breakthrough (n=12) | 2 × Vac + BA.2 breakthrough (n=12) | 3 × Vac + BA.2 breakthrough (n=12) | 2 × Vac + BA.5 breakthrough (n=9) | 3 × Vac + BA.5 breakthrough (n=11) | 3 × Vac + BF.7 breakthrough (n=12) |
|---|---|---|---|---|---|---|---|---|
| Age(years), median (range) | 44.3(28-63) | 39.7(20-53) | 47(35-55) | 43(25-66) | 37(32-66) | 30.8(23-47) | 30.6(22-47) | 40.1 (34-50) |
| Male, n (%) | 7(63.6%) | 3(27.2%) | 5(41.6%) | 9 (75.0%) | 8(66.7%) | 6(66.6%) | 5(45.5%) | 4(33.3%) |
| BMI (kg/m$^2$), mean (SD) | 24.7(3.1) | 23.9(3.7) | 29(10.7) | 25.1(2.6) | 25.2(3.2) | 24(3.2) | 20.9(2.2) | 23.9(3.3) |
| Breakthrough infections days after the last Coronavirus vaccines, median (range) | N/A | N/A | 76.4 (34-128) | ND | ND | 458.3 (325-587) | 313.5 (220-387) | 407.4 (363-432) |
| Serum samples collection days after the vaccination or infection, median (range) | 17.5（15-22） | 18（18） | 156.6（153-161） | 180(164-202) | 176(167-196) | 25.2（21-34） | 21.9（18-24） | 14.1（10-19） |
| Comorbidities (%) | | | | | | | | |
| Any, n (%) | 0(0%) | 0(0%) | 2(16.6%) | 3(25%) | 0(0%) | 0(0%) | 0(0%) | 0(0%) |
| HTN, n (%) | 0(0%) | 0(0%) | 2(16.6%) | 3(25%) | 0(0%) | 0(0%) | 0(0%) | 0(0%) |
| CAD, n (%) | 0(0%) | 0(0%) | 0(0%) | 0(0%) | 0(0%) | 0(0%) | 0(0%) | 0(0%) |
| DM, n (%) | 0(0%) | 0(0%) | 0(0%) | 0(0%) | 0(0%) | 0(0%) | 0(0%) | 0(0%) |
| NASH, n (%) | 0(0%) | 0(0%) | 0(0%) | 0(0%) | 0(0%) | 0(0%) | 0(0%) | 0(0%) |
| Arrhy, n (%) | 0(0%) | 0(0%) | 0(0%) | 0(0%) | 0(0%) | 0(0%) | 0(0%) | 0(0%) |
| Asthma, n (%) | 0(0%) | 0(0%) | 0(0%) | 0(0%) | 0(0%) | 0(0%) | 0(0%) | 0(0%) |
| Rhinitis, n (%) | 0(0%) | 0(0%) | 0(0%) | 0(0%) | 0(0%) | 0(0%) | 0(0%) | 0(0%) |
| Urticaria, n (%) | 0(0%) | 0(0%) | 0(0%) | 0(0%) | 0(0%) | 0(0%) | 0(0%) | 0(0%) |

N/A, not applicable. ND, no data. BMI, body mass index. CAD, coronary artery disease. HTN, hypertension. DM, diabetes mellitus. Arrhy, arrhythmia, NASH, non-alcoholic steatohepatitis.
